# A three dimensional multiplane kinematic model for bilateral hind limb gait analysis in cats

**DOI:** 10.1101/320812

**Authors:** Nathan P. Brown, Gina E. Bertocci, Kimberly A. Cheffer, Dena R. Howland

**Affiliations:** Department of Bioengineering, J.B. Speed School of Engineering, University of Louisville, Louisville, Kentucky, United States of America; Kentucky Spinal Cord Injury Research Center, University of Louisville, Louisville, Kentucky, United States of America; Research Service, Robley Rex VA Medical Center, Louisville, Kentucky, United States of America; Department of Anatomical Sciences and Neurobiology, School of Medicine, University of Louisville, Louisville, Kentucky, United States of America; Department of Neurological Surgery, School of Medicine, University of Louisville, Louisville, Kentucky, United States of America

## Abstract

Background: Kinematic gait analysis is an important noninvasive technique used for quantitative evaluation and description of locomotion and other movements in healthy and injured populations. Three dimensional (3D) kinematic analysis offers additional outcome measures including internal-external rotation not characterized using sagittal plane analysis techniques.

Methods: The objectives of this study were to 1) develop and evaluate a 3D hind limb multiplane kinematic model for gait analysis in cats using joint coordinate systems, 2) implement and compare two 3D stifle (knee) prediction techniques, and 3) compare flexion-extension determined using the multiplane model to a sagittal plane model. Walking gait was recorded in 3 female adult cats (age = 2.9 years, weight = 3.5 ± 0.2 kg). Kinematic outcomes included flexion-extension, internal-external rotation, and abduction-adduction of the hip, stifle, and tarsal (ankle) joints.

Results: Each multiplane stifle prediction technique yielded similar findings. Joint angles determined using markers placed on skin above bony landmarks *in vivo* were similar to joint angles determined using a feline hind limb skeleton in which markers were placed directly on landmarks *ex vivo*. Differences in hip, stifle, and tarsal joint flexion-extension were demonstrated when comparing the multiplane model to the sagittal plane model.

Conclusions: This multiplane cat kinematic model can predict joint rotational kinematics as a tool that can quantify frontal, transverse, and sagittal plane motion. This model has multiple advantages given its ability to characterize joint internal-external rotation and abduction-adduction. A further, important benefit is greater accuracy in representing joint flexion-extension movements.

## Introduction

Kinematic gait analysis is an important noninvasive technique that is used to quantify locomotion and other movements in healthy and injured populations [1–3]. Most animal model research has used sagittal plane kinematic analysis techniques to evaluate changes in locomotor features across a variety of gait tasks due to injury and/or recovery [4–10]. A significant drawback to sagittal plane kinematic analyses is the inability to quantify angular changes associated with out-of-plane motions such as internal-external rotation or abduction-adduction typical of normal gait and exaggerated in pathological gait following a wide variety of neurodegenerative disorders [3, 11, 12]. Sagittal plane kinematic models use either 2D or 3D marker coordinate data. Using 2D coordinate data constrains all marker coordinates to a static global sagittal plane that does not change regardless of direction of movement. In contrast, 3D coordinate data dynamically defines a local sagittal plane that contains the surface-mounted markers corresponding to the segments comprising the joint of interest [13, 14]. Given that most joints have three rotational degrees of freedom (flexion-extension, internal-external rotation, and abduction-adduction), both sagittal plane techniques are limiting as they capture only a single representational movement (flexion-extension) and cannot account for rotation into or out of the sagittal plane (i.e. internal-external rotation). Joint flexion-extension angles determined using sagittal plane kinematic models actually represent a composite of sagittal, transverse, and frontal plane movements. Recently, a composite sagittal plane-frontal plane technique was developed [13]. Fore and hind limb frontal plane kinematics were described as the angle between a line connecting the hip joints and shoulder joints, respectively, and the sagittal plane defined by the fore and hind limbs, respectively [13, 14]. Rotational components in the transverse plane (i.e. internal-external rotation), however, are not quantified using this technique, and the knee location in this model may be influenced by skin motion artifact. A multiplane model has the potential to capture greater detail on clinically relevant changes in kinematics following neural damage and recovery and allows an improved understanding of kinematic motions beyond that which can be determined using sagittal plane and composite sagittal plane-frontal plane techniques.

Gait analysis using multiplane kinematics allows for quantification of joint angles in the sagittal, transverse, and frontal planes to describe flexion-extension, internal-external rotation, and abduction-adduction [15–17]. The joint coordinate system approach utilizes rotations between segment-embedded, anatomically-oriented coordinate systems based on bony landmarks to describe clinically relevant 3D joint motions [15]. Multiplane kinematic models have been implemented in humans [15, 18, 19], dogs [17, 20], and horses [16, 21, 22]. Rat and cat hind limb musculoskeletal anatomy and hind limb coordinate systems have been described in detail previously based on anatomical axes [13, 23, 24], and progress has been made towards development of a multiplane kinematic model in cats [13, 14]. The cat neurologic model has contributed significantly to our understanding of motor control [6–8, 10, 25–30]. Therefore, 3D kinematic analyses will enhance quantification of neurologic deficits and functional recovery. A multiplane kinematic model to describe sagittal, frontal, and transverse joint rotations during gait using skin surface mounted motion capture markers in cats, however, has not been developed. Similar to rodent gait analysis marker sets [31, 32], a cat marker set used to define joint coordinate systems must account for skin motion at the stifle (knee) which may lead to unreliable kinematic analysis [33]. Previous studies evaluating sagittal plane kinematics have defined the stifle using a predicted marker based on a vector projection of the tibia-fibula segment length in cats [25, 26, 30], femur and tibia segment lengths in cats [34], and femur and tibia segment lengths in rats [31, 32]. Stifle prediction methods demonstrated superiority to using a stifle skin marker in the analysis in rats when compared to x-ray cinematography [31].

To our knowledge no study has investigated sagittal, frontal, and transverse plane kinematics of walking gait in the cat, and a robust pelvis and hind limb kinematic model has not been described. The purpose of this study was to 1) develop and evaluate a 3D multiplane kinematic model of the cat pelvis and hind limbs for gait analysis using anatomically-oriented joint coordinate systems, 2) implement and compare two 3D stifle prediction techniques, and 3) compare flexion-extension determined using the 3D multiplane model to a sagittal plane model.

## Materials and methods

### Kinematic gait trials

Four purpose-bred specific-pathogen free spayed female adult cats were included in this study. All procedures were approved by the University of Louisville and Robley Rex VA Institutional Animal Care and Use Committees. Three cats were conditioned to cross a 4.6 m runway (0.36 m width) at their self-selected walking speeds approximately 30 times daily, 5 times per week for food rewards. Motion capture data were acquired using a 3D test space encompassing the entire width and length of the walkway. Data were collected at 100 Hz using 10 infrared motion capture tracking cameras (4 MX T160, Vicon, Centennial, CO; 6 MX T40-S, Vicon, Centennial, CO), and left and right sagittal views were recorded at 100 Hz using synched videography (2 Bonita 720c, Vicon, Centennial, CO, and 2 HDR-CX220, Sony, Tokyo, Japan). A minimum of 3 motion capture cameras were required to reconstruct each motion capture marker. Fur was shaved, and reflective motion capture markers (6.4 mm diameter) were placed bilaterally on the skin over bony landmarks of the pelvis and hind limbs. Bilateral marker locations included the iliac crest, ischial tuberosity, greater trochanter, lateral malleolus, medial malleolus, caudolateral aspect of the calcaneus, laterodistal aspect of the 5^th^ metatarsal, and mediodistal aspect of the 2^nd^ metatarsal. An additional marker was placed bilaterally approximately 2.5 cm proximal to the lateral malleolus on the lateral aspect of the fibula. Three representative overground gait trials with a minimum of 3 consecutive step cycles each at consistent speeds were selected for each subject. Each step cycle was resampled into percent step cycle and divided into stance and swing phases. Gaps in data < 5 frames were filled using Woltring quintic splines. The fourth cat was included for model validation, was not a subject during *in vivo* data collection, and was euthanized for reasons unrelated to this study. For use during model validation, the pelvis and hind limbs were disarticulated at the sacrum, and all soft tissues were excised except hip, stifle, and tarsal joint connective tissues (i.e., joint capsule and ligaments) which were left intact. The specimen initially was exposed to a fixative (4% buffered paraformaldehyde) and then maintained in water at 4°C to prevent specimen deterioration and stiffening of the connective tissues.

### Prediction of stifle

Due to skin motion artifact across the stifle joint, a retroreflective marker was not placed on the cat stifle. Instead, two techniques were used to determine a lateral stifle virtual marker. For the first technique (unadjusted tibia axis technique), the lateral stifle virtual marker was defined as the midpoint of the line connecting 1) the vector of length equal to the fibula length originating at the lateral malleolus projected through the vector marker and 2) the vector of length equal to the femur length originating at the greater trochanter projected towards the fibula vector endpoint (Fig 1). For the second technique (adjusted tibia axis technique), the lateral stifle virtual marker was determined as described in the first technique, but the tibia anatomical axes were redefined using the calculated lateral stifle virtual marker instead of the vector marker (Fig 1, Table 1). The femur and fibula lengths for each subject were determined by palpating bony landmarks and measuring the distances from the greater trochanter to the distal aspect of the femoral condyle and the lateral malleolus to the fibular head, respectively.

**Fig 1.**
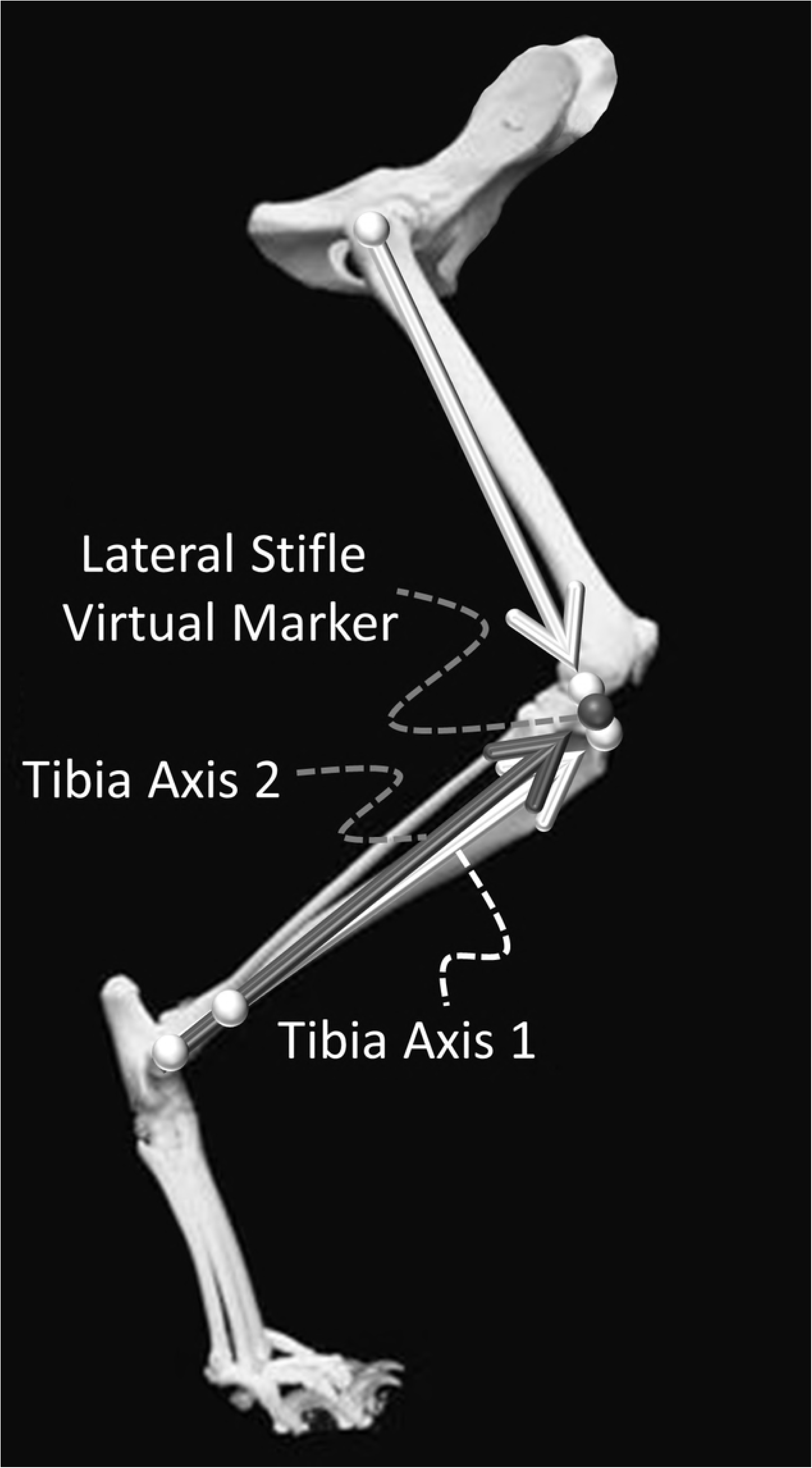
Lateral stifle virtual marker (dark sphere) projection techniques. Tibia axis 1 (light arrow along the tibia) was defined as the vector of length equal to the fibula length originating at the lateral malleolus projected through the vector marker. The femoral axis was defined as the vector of length equal to the femur length originating at the greater trochanter projected towards the fibula vector endpoint. The lateral stifle virtual marker was defined as the midpoint of the line connecting the endpoints of tibia axis 1 and the femoral axis. Tibia axis 2 (dark arrow along the tibia) was defined as the vector of length equal to the fibular length originating at the lateral malleolus projected towards the lateral stifle virtual marker determined previously.

**Table 1.** Cat model anatomical coordinate systems and directional unit vector axes.

| Coordinate system | Origin | Z axis | X axis | Y axis |
| --- | --- | --- | --- | --- |
| Pelvis | RIC | $\hat{z} = \frac{\bar{V}_{RIC} - \bar{V}_{LIC}}{\vert \bar{V}_{RIC} - \bar{V}_{LIC}\vert }$ | $\hat{x} = \frac{(\bar{V}_{RIC} - \bar{V}_{RIT}) \times \hat{z}}{\vert (\bar{V}_{RIC} - \bar{V}_{RIT}) \times \hat{z}\vert }$ | $\hat{y} = \hat{z} \times \hat{x}$ |
| Right Femur | RGT | $\hat{z} = \hat{x} \times \hat{y}$ | $\hat{x} = \frac{(\bar{V}_{RMM} - \bar{V}_{RLM}) \times \hat{y}}{\vert (\bar{V}_{RMM} - \bar{V}_{RLM}) \times \hat{y}\vert }$ | $\hat{y} = \frac{\bar{V}_{RGT} - \bar{V}_{RS}}{\vert \bar{V}_{RGT} - \bar{V}_{RS}\vert }$ |
| Right Tibia<sub>1</sub> | RLM | $\hat{z} = \frac{\bar{V}_{RLM} - \bar{V}_{RMM}}{\vert \bar{V}_{RLM} - \bar{V}_{RMM}\vert }$ | $\hat{x} = \frac{(\bar{V}_{RV} - \bar{V}_{RLM}) \times \hat{z}}{\vert (\bar{V}_{RV} - \bar{V}_{RLM}) \times \hat{z}\vert }$ | $\hat{y} = \hat{z} \times \hat{x}$ |
| Right Tibia<sub>2</sub> | RLM | $\hat{z} = \frac{\bar{V}_{RLM} - \bar{V}_{RMM}}{\vert \bar{V}_{RLM} - \bar{V}_{RMM}\vert }$ | $\hat{x} = \frac{(\bar{V}_{RS} - \bar{V}_{RLM}) \times \hat{z}}{\vert (\bar{V}_{RS} - \bar{V}_{RLM}) \times \hat{z}\vert }$ | $\hat{y} = \hat{z} \times \hat{x}$ |
| Right Tarsus | RC | $\hat{z} = \frac{\bar{V}_{R5MT} - \bar{V}_{R2MT}}{\vert \bar{V}_{R5MT} - \bar{V}_{R2MT}\vert }$ | $\hat{x} = \frac{(\bar{V}_{RC} - \bar{V}_{R5MT}) \times \hat{z}}{\vert (\bar{V}_{RC} - \bar{V}_{R5MT}) \times \hat{z}\vert }$ | $\hat{y} = \hat{z} \times \hat{x}$ |
| Left Femur | LGT | $\hat{z} = \hat{x} \times \hat{y}$ | $\hat{x} = \frac{(\bar{V}_{LLM} - \bar{V}_{LMM}) \times \hat{y}}{\vert (\bar{V}_{LLM} - \bar{V}_{LMM}) \times \hat{y}\vert }$ | $\hat{y} = \frac{\bar{V}_{LGT} - \bar{V}_{LS}}{\vert \bar{V}_{LGT} - \bar{V}_{LS}\vert }$ |
| Left Tibia<sub>1</sub> | LLM | $\hat{z} = \frac{\bar{V}_{LMM} - \bar{V}_{LLM}}{\vert \bar{V}_{LMM} - \bar{V}_{LLM}\vert }$ | $\hat{x} = \frac{(\bar{V}_{LV} - \bar{V}_{LLM}) \times \hat{z}}{\vert (\bar{V}_{LV} - \bar{V}_{LLM}) \times \hat{z}\vert }$ | $\hat{y} = \hat{z} \times \hat{x}$ |
| Left Tibia<sub>2</sub> | LLM | $\hat{z} = \frac{\bar{V}_{LMM} - \bar{V}_{LLM}}{\vert \bar{V}_{LMM} - \bar{V}_{LLM}\vert }$ | $\hat{x} = \frac{(\bar{V}_{LS} - \bar{V}_{LLM}) \times \hat{z}}{\vert (\bar{V}_{LS} - \bar{V}_{LLM}) \times \hat{z}\vert }$ | $\hat{y} = \hat{z} \times \hat{x}$ |
| Left Tarsus | LC | $\hat{z} = \frac{\bar{V}_{L2MT} - \bar{V}_{L5MT}}{\vert \bar{V}_{L2MT} - \bar{V}_{L5MT}\vert }$ | $\hat{x} = \frac{(\bar{V}_{LC} - \bar{V}_{L5MT}) \times \hat{z}}{\vert (\bar{V}_{LC} - \bar{V}_{L5MT}) \times \hat{z}\vert }$ | $\hat{y} = \hat{z} \times \hat{x}$ |

V = vector, RIC = right iliac crest, RIT = right ischial tuberosity, RGT = right greater trochanter, RS = right stifle, RLM = right lateral malleolus, RMM = right medial malleolus, RV = right vector, RC = right calcaneus, R5MT = right 5<sup>th</sup> metatarsal, R2MT = right 2<sup>nd</sup> metatarsal, LIC = left iliac crest, LGT = left greater trochanter, LS = left stifle, LLM = left lateral malleolus, LMM = left medial malleolus, LV = left vector, LC = left calcaneus, L5MT = left 5<sup>th</sup> metatarsal, L2MT = left 2<sup>nd</sup> metatarsal

Left and right tibial coordinate systems were determined using two techniques and are identified with subscript numerals.

### Pelvis and bilateral hind limb anatomical coordinate systems

Anatomical segment coordinate systems were developed for the pelvis and bilateral femur, tibia, and tarsus based on markers placed on the skin (Fig 2). Each anatomical segment coordinate system defined clinically relevant feline pelvis and hind limb anatomical axes adapted from humans [15] and canines [17, 20, 35] based upon the International Society of Biomechanics recommendations [18, 36]. Coordinate axes directions and the coordinate system origins for each segment are listed in Table 1.

**Fig 2.**
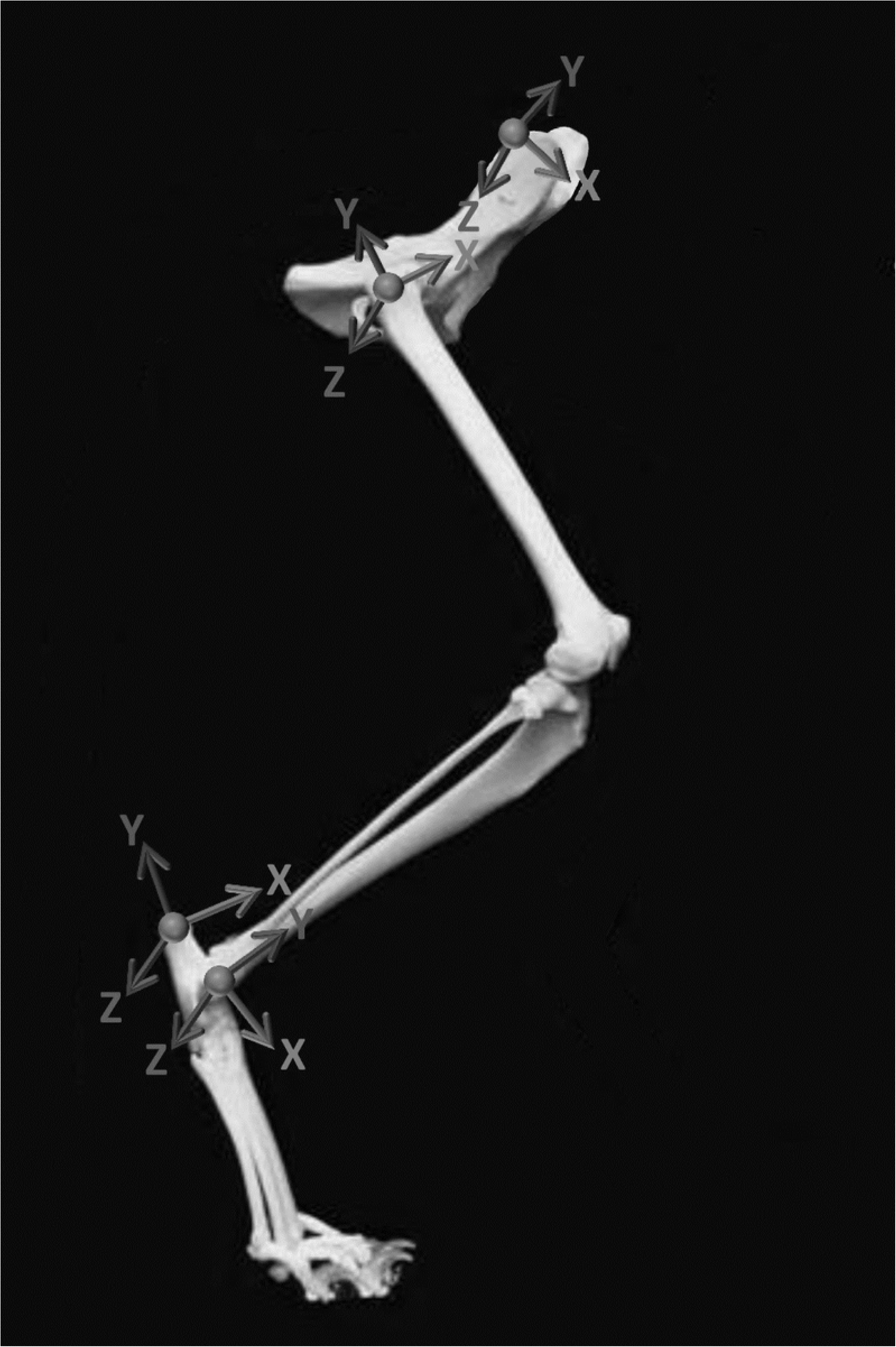
Cat model anatomical coordinate systems. Anatomical coordinate systems were determined for the pelvis, femur, tibia, and tarsus.

### Joint coordinate systems

Rotations of the distal segment relative to the proximal segment about three clinical axes were determined using joint coordinate systems [15, 17]. Flexion-extension was defined as rotation about a unit vector along the proximal segment z-axis, internal-external rotation was defined as rotation about a unit vector along the distal segment y-axis, and abduction-adduction was defined as rotation about a unit vector that is perpendicular to both axes.

### Kinematic analysis

Data were analyzed using motion analysis software (Nexus 2.1.1, Vicon, Centennial, CO) and a custom script (BodyLanguage, Vicon, Centennial, CO). Three-dimensional kinematics were determined for the hip, stifle, and tarsal joints of each hind limb. Kinematic angles in the sagittal (flexion-extension), transverse (internal-external rotation), and frontal (abduction-adduction) planes were determined. Stifle kinematic analysis was conducted for each stifle prediction technique. Flexion-extension determined using the multiplane kinematic model was compared to flexion-extension determined using a 3 coordinate sagittal plane kinematic model [30]. For the sagittal plane model, flexion-extension of the hip, stifle, and tarsal joints was calculated using the iliac crest, greater trochanter, lateral stifle virtual marker, lateral malleolus, and 5^th^ metatarsal markers. Kinematic outcomes were verified using synched video analysis.

### Model validation

The multiplane kinematic model was applied to the right hind limb of the cadaveric specimen. During testing the specimen pelvis was rigidly fixed using two clamps attached to the right iliac wing and ischial spine at the obturator foramen. The hip, stifle, and tarsal joints could move freely within the constraints of the connective tissues. Markers were adhered directly to bony locations previously described for placement of skin markers. Additionally, a marker triad representing bone oriented coordinate systems was adhered to each hind limb segment (pelvis, femur, tibia-fibula, and tarsus) (Fig 3). Each triad consisted of a triangular flat acrylic glass surface (25 mm × 25 mm × 3 mm) and 3 markers: an origin marker and two markers placed 19 mm in the positive X and Y directions. The femur, tibia-fibula, and tarsus triad X axes were aligned caudal to cranial, and the triad Y axes were aligned distal to proximal. The pelvis triad X axis was aligned medial to lateral, and the triad Y axis was aligned caudal to rostral. Each marker triad Z axis was the cross product of the X and Y axes. The femur, tibia-fibula, and tarsus triads were placed laterally approximately midway from the proximal and distal aspects. The pelvis triad was placed between the iliac wings and dorsal to the pelvis using a 51-mm extension to prevent interference with the pelvis clamps.

**Fig 3.**
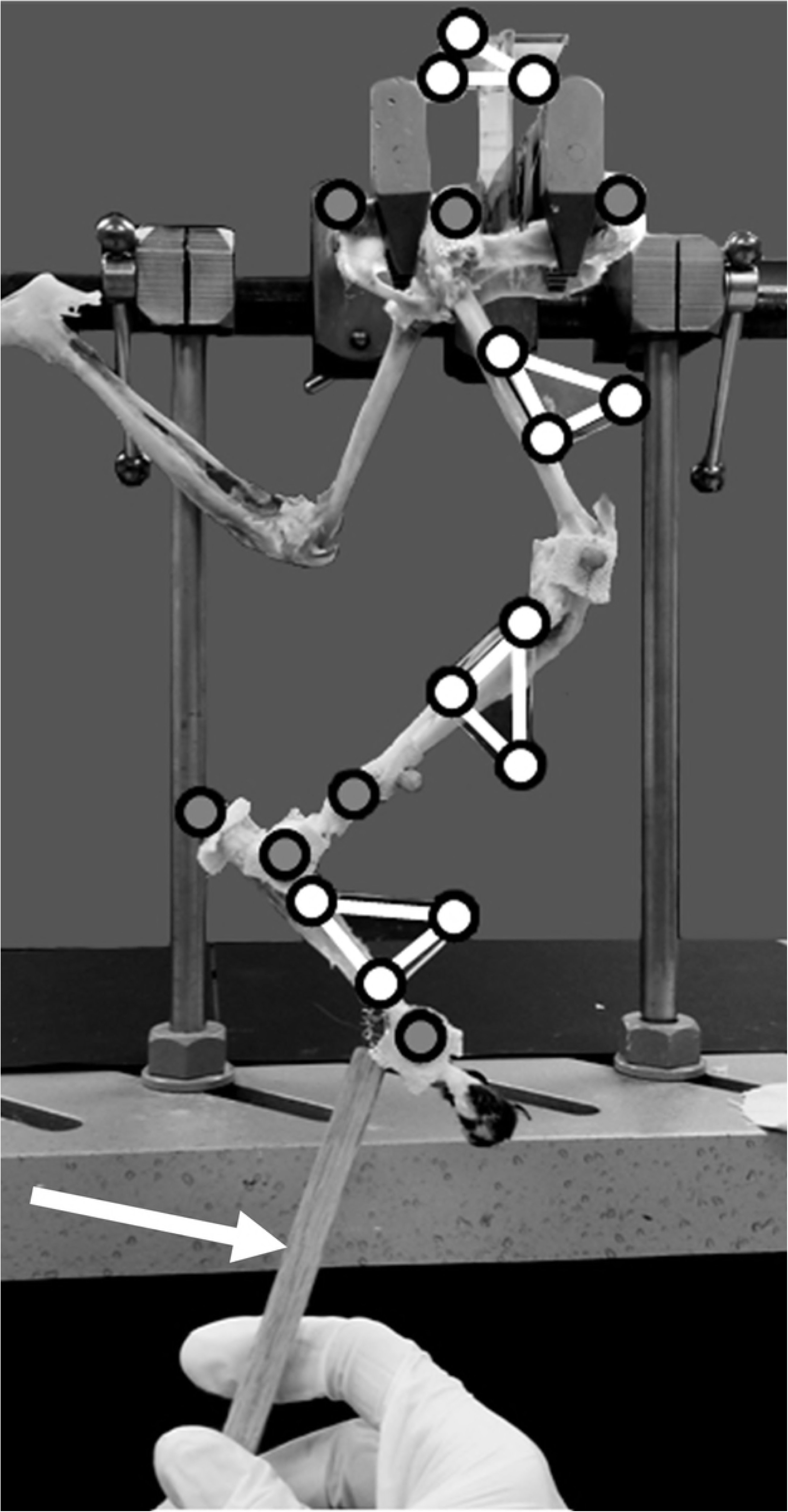
Hind limb specimen with bony landmark markers (gray circles) and marker triads (white circles at triangle vertices). Markers and marker triads are adhered along the pelvis and a single limb. The left hind limb was suspended to prevent obstruction of the right hind limb markers. The right hind limb was manipulated using the rigid extension (identified with a white arrow) attached to the metatarsal bones distal to the tarsus triad. Reflective markers can be seen at the stifle and anterior tibia, but these markers were not used in the current study.

The hind limb specimen was manually moved through a walking gait motion (metatarsus moved elliptically in the sagittal plane to mimic walking). Hip, stifle, and tarsal joints were manipulated collectively in 3D space using a rigid wooden extension attached to the metatarsus. Thin metal wire was threaded through the metatarsus connective tissue and secured to an eyebolt affixed to the rigid extension. The rigid extension could rotate relative to the tarsus while providing translational constraint. Nine cycles were recorded, and data were normalized by resampling into percent cycle.

The joint coordinate system was used to determine 3D joint angles using the marker triads on each segment. Joint angles determined using the bony landmark marker set and the marker triads were compared. The bony landmark marker set and the marker triad marker set defined coordinate system orientations on each segment. Therefore, small differences in marker placement and coordinate system alignment were present and unavoidable. Offset in coordinate system orientations defined by the bony landmark marker set and the marker triad on each segment due to alignment differences between the marker sets was eliminated using coordinate transformation. The bony landmark coordinate system joint angles were related to the marker triad coordinate system joint angles using the following transformation at each joint:

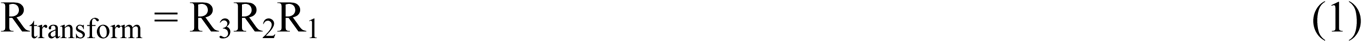

where R_3_ is the rotation matrix from the proximal segment bony landmark coordinate system to the proximal segment marker triad coordinate system, R_2_ is the rotation matrix from the distal segment bony landmark coordinate system to the proximal segment bony landmark coordinate system, and R_1_ is the rotation matrix from the distal segment marker triad coordinate system to the distal segment bony landmark coordinate system. The rotation matrices R_1_, R_2_, and R_3_, are each notationally equivalent to

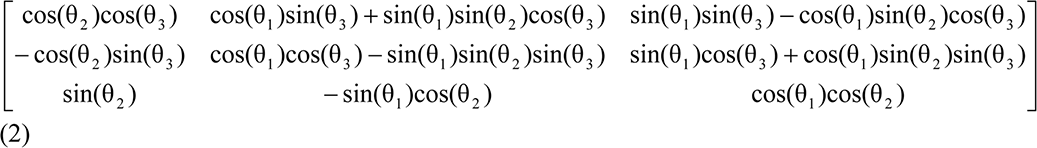

where θ_1_, θ_2_, and θ_3_ are the three rotations to transform one coordinate system into another coordinate system [37]. R_1_ and R_3_ were determined for one frame with the hind limb mounted in a neutral position (i.e., the limb was not subjected to forces using the rigid wooden extension) and then applied to all frames. R_2_ corresponds to the joint angles determined at each frame using the bony landmark coordinate systems of the proximal and distal segments. R_transform_ joint angles were compared to joint angles determined using the marker triad coordinate systems of the proximal and distal segments.

## Results

Ages were 2.9 years, mean weight was 3.5 ± 0.2 kg, and mean walking speed was 0.73 ± 0.08 m/s for the three cats performing gait trials. Age was 1.4 years and weight was 3.4 kg for the cat used for model validation.

### Stifle prediction comparison

Multiplane stifle kinematics determined using both prediction techniques were similar, and a representative hind limb is shown in Fig 4.

**Fig 4.**
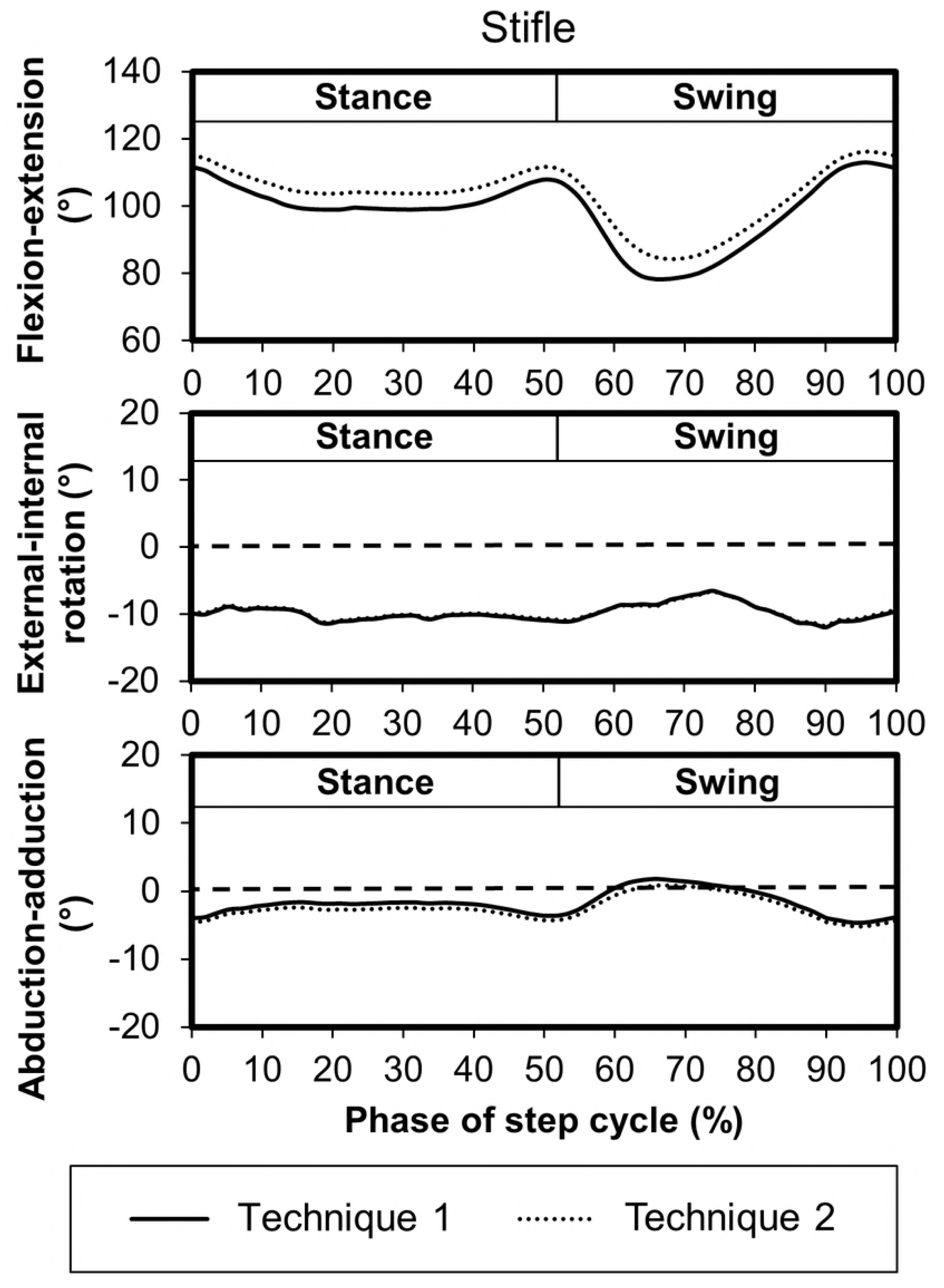
Mean stifle joint kinematics for a representative hind limb using both stifle prediction techniques during the step cycle for gait. Technique 1 corresponds to the unadjusted tibia axis, and technique 2 corresponds to the adjusted tibia axis. Flexion-extension angles correspond to the absolute angle between the segments while positive values indicate external rotation and abduction, and negative values correspond to internal rotation and adduction. The dashed horizontal lines indicate 0° (neutral) on external-internal rotation and abduction-adduction graphs.

### Kinematics

Bilateral 3D kinematics for the hip, stifle, and tarsal joints during stance and swing were determined, and a representative hind limb is shown in Fig 5. Mean and SD joint angles, maximum joint angles, minimum joint angles, and range of motion throughout the step cycle for each joint rotation were compared (Table 2). Data reported in Fig 5 and Table 2 are for the same cat. Consistent with what is known, these data show that the hip joint extends throughout stance, reaching maximum extension at the end of stance (transition from weight support to non-weight support), and then flexes throughout swing to place the paw again onto the walking surface at the beginning of stance to accept weight. The stifle joint remains extended throughout stance, flexes in early swing to maintain the paw above ground, and extends in late swing to place the paw on the ground. The tarsal joint flexes slightly in early stance, remains extended throughout stance, flexes in early swing to maintain the paw above ground, and extends in late swing to place the paw on the ground. These data also support that prominent changes in rotation and abduction-adduction often occur at transitions between stance and swing. For instance, for the cat depicted in Fig 5, hip external rotation increases in early stance to facilitate forward motion of the pelvis relative to the femur. Furthermore, during early swing external rotation in the tarsal joint increases allowing clearance of the contralateral limb, and external rotation decreases prior to stance for paw placement. Stifle and tarsal internal-external rotation and abduction-adduction remained relatively unchanged during stance providing limb stability during weight bearing while the contralateral limb was in swing.

**Fig 5.**
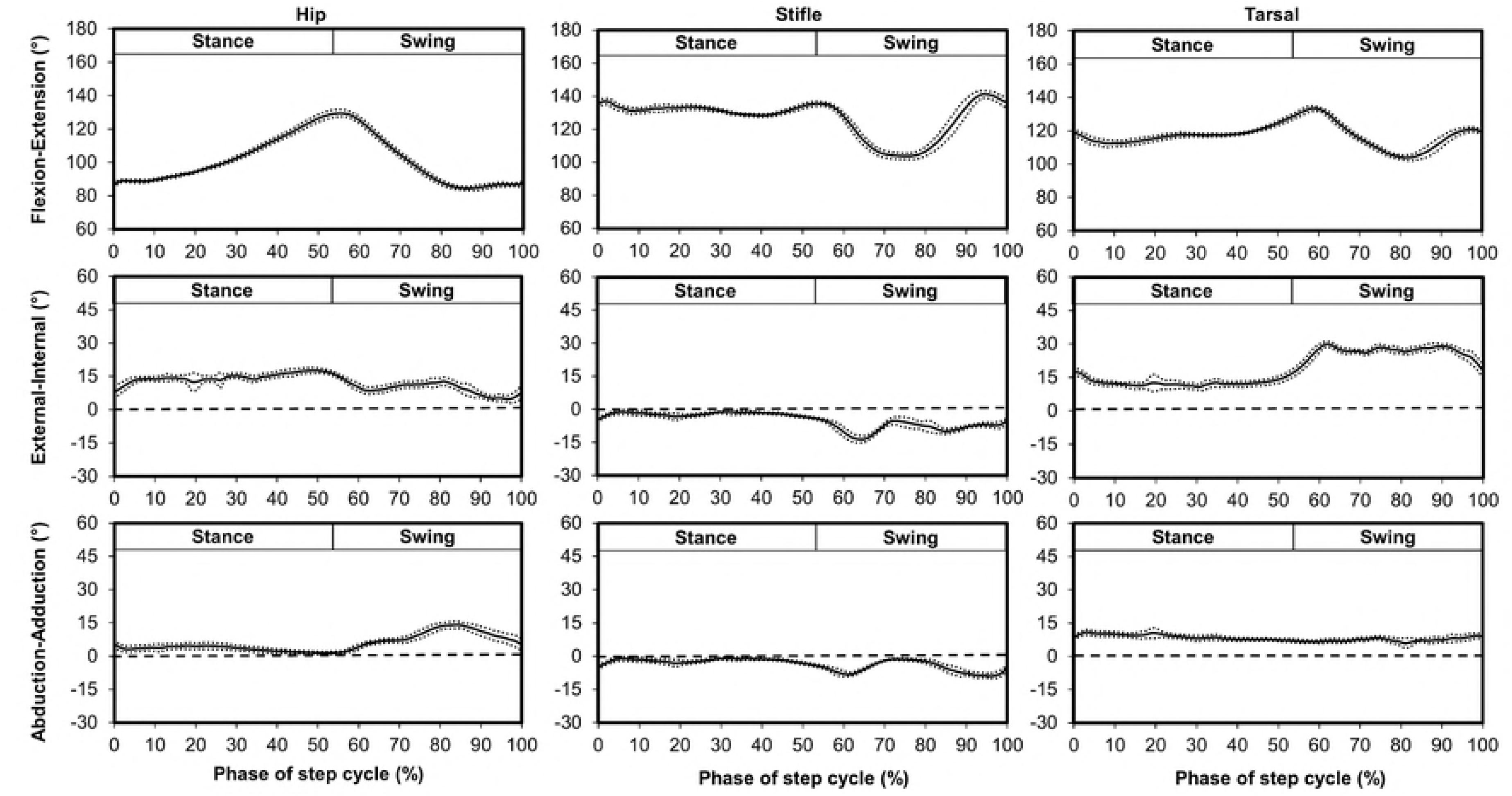
Mean (solid line) ± SD (dotted line) three dimensional joint coordinate system kinematics for the hip, stifle, and tarsal joints for a representative hind limb during the step cycle for gait. Flexion-extension angles correspond to the absolute angle between the segments while external rotation and abduction correspond to positive and internal rotation and adduction correspond to negative. The dashed horizontal lines indicate 0° on external-internal rotation and abduction-adduction graphs.

**Table 2.** Bilateral mean and SD, maximum, and minimum joint angles, and range of motion (ROM) throughout the average step cycle for each joint rotation from a representative cat using the multiplane model.

|  | Mean (SD) |  | Max | Min | ROM |
| --- | --- | --- | --- | --- | --- |
| **Left hip joint (°)** |  |  |  |  |  |
| Flexion-extension | 101.7 | (15.7) | 129.6 | 82.2 | 47.4 |
| Internal-external rotation | 4.9 | (3.9) | 10.9 | -3.7 | 14.6 |
| Abduction-adduction | -7.0 | (4.6) | 0.5 | -14.4 | 14.9 |
| **Left stifle joint (°)** |  |  |  |  |  |
| Flexion-extension | 128.1 | (10.9) | 143.7 | 106.5 | 37.2 |
| Internal-external rotation | -8.6 | (2.8) | -3.2 | -14.5 | 11.3 |
| Abduction-adduction | -7.4 | (3.5) | -0.9 | -16.2 | 15.3 |
| **Left tarsal joint (°)** |  |  |  |  |  |
| Flexion-extension | 117.0 | (6.0) | 129.8 | 105.8 | 24.0 |
| Internal-external rotation | 9.3 | (8.2) | 24.7 | -0.1 | 24.9 |
| Abduction-adduction | 5.3 | (2.3) | 9.4 | 0.7 | 8.7 |
| **Right hip joint (°)** |  |  |  |  |  |
| Flexion-extension | 101.7 | (14.8) | 129.5 | 84.5 | 45.1 |
| Internal-external rotation | 12.3 | (3.4) | 17.7 | 4.8 | 12.9 |
| Abduction-adduction | 5.7 | (3.7) | 14.1 | 1.4 | 12.7 |
| **Right stifle joint (°)** |  |  |  |  |  |
| Flexion-extension | 126.9 | (11.0) | 141.2 | 103.7 | 37.5 |
| Internal-external rotation | -5.0 | (3.5) | -1.1 | -13.5 | 12.4 |
| Abduction-adduction | -3.5 | (2.5) | -0.9 | -8.9 | 8.0 |
| **Right tarsal joint (°)** |  |  |  |  |  |
| Flexion-extension | 117.2 | (7.1) | 133.4 | 103.8 | 29.6 |
| Internal-external rotation | 19.0 | (7.2) | 29.9 | 19.3 | 10.7 |
| Abduction-adduction | 8.2 | (1.2) | 10.9 | 5.9 | 5.0 |

Flexion-extension angles correspond to the absolute angle between the segments while external rotation and abduction correspond to positive and internal rotation and adduction correspond to negative.

**Fig 6.**
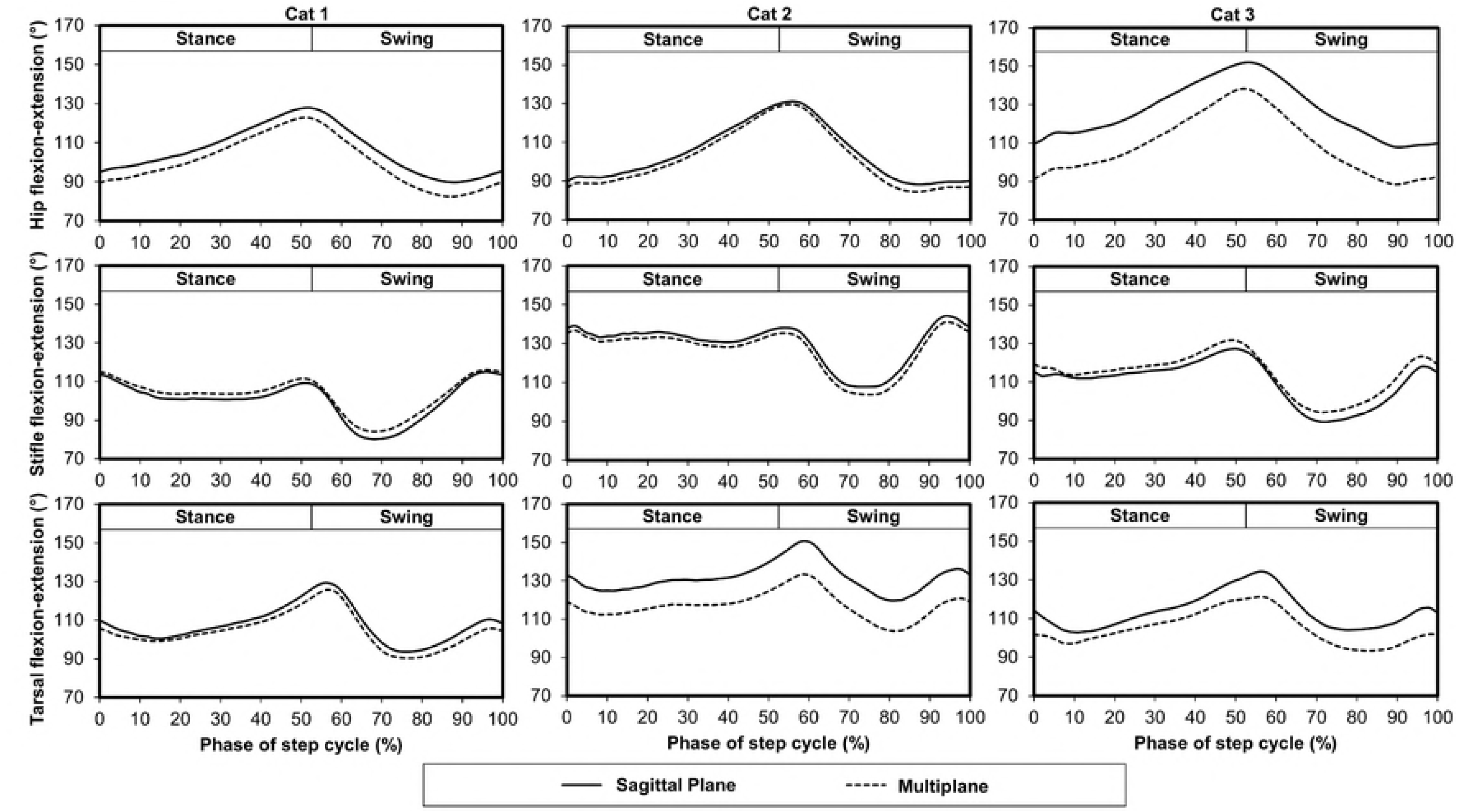
Mean flexion-extension for the hip, stifle, and tarsal joints for a representative limb from each cat using the multiplane and sagittal plane kinematic models during the step cycle for gait. Cat 1 corresponds to the cat in Figure 4 and Cat 2 corresponds to the cat in Figure 5.

**Fig 7.**
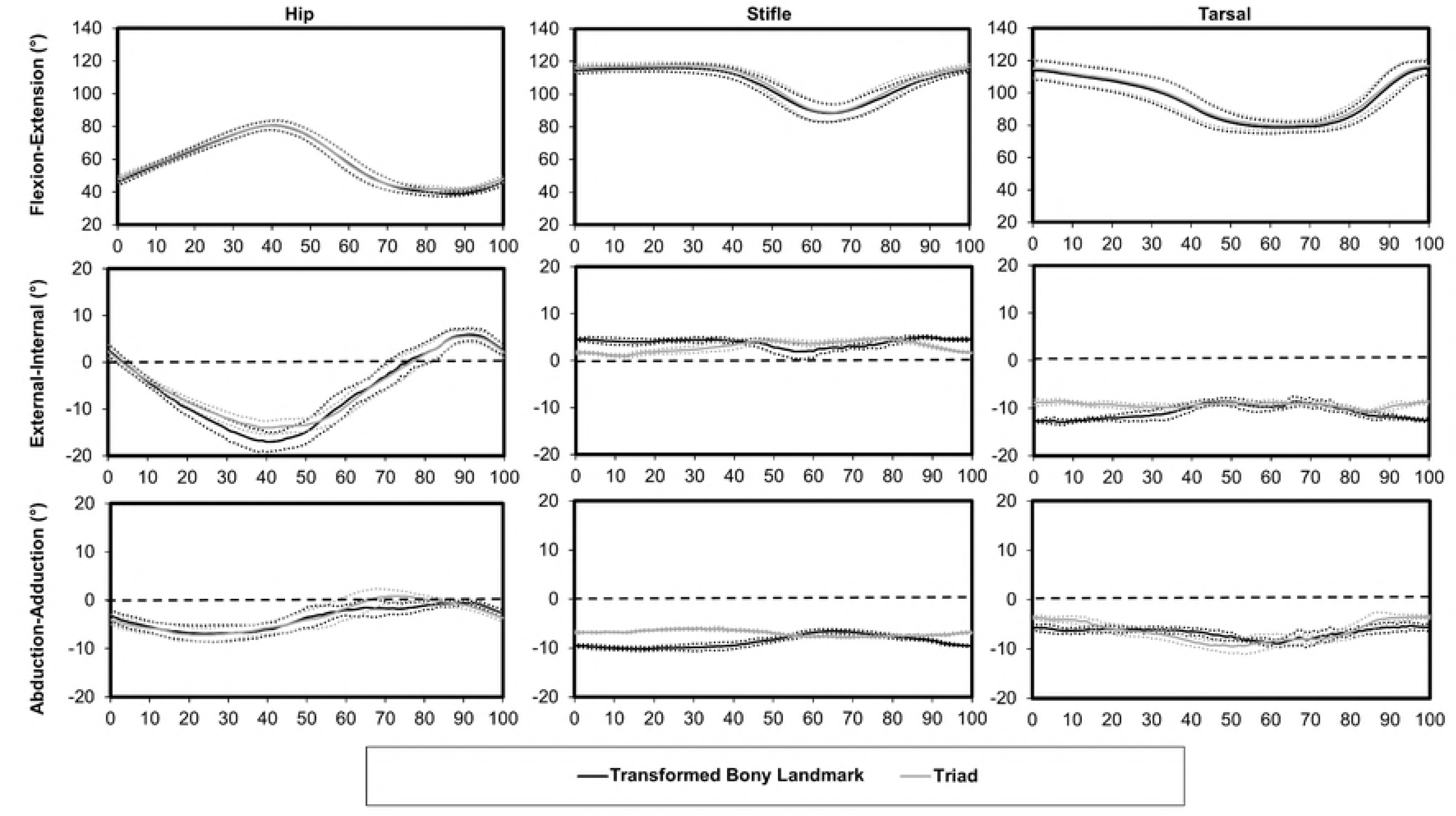
Hip, stifle, and tarsal mean (solid line) ± SD (dotted line) kinematics for the *ex vivo* walking task using the transformed bony landmark marker set and the triad marker set. The horizontal dashed line indicates 0° on external-internal rotation and abduction-adduction graphs.

## Discussion

A feline bilateral hind limb multiplane kinematics model was successfully developed, applied, and tested during walking gait. This multiplane model will allow quantification of hind limb flexion-extension, internal-external rotation, and abduction-adduction. We anticipate that this model will serve as an effective tool to compare gait characteristics across a variety of walking tasks and pathologic conditions and enhance our understanding of biomechanical changes associated with neurological conditions. The cat model has been important for studies which contribute to our current understanding of motor control [6–8, 10, 25–30]. Using a multiplane model in this species will provide opportunity for additional insights such as internal-external rotation not feasible with current, standard kinematic models used for quadrupeds.

### 3D cat hind limb kinematics

Rotation is an important component of limb movement during key sub-phases [20] and can be captured using a multiplane kinematic approach. Although sagittal plane and multiplane kinematics models have demonstrated general agreement in flexion-extension [38], multiplane kinematics models provide additional information including internal-external rotation that more fully characterize gait by identifying additional features critical in differentiating healthy and pathologic movements [20]. For example, internal rotation of the tibia increases in dogs with cranial cruciate ligament injury [39, 40], and hip internal rotation increases in humans with a crouch gait.[2] Flexion-extension waveforms reported in the current study demonstrate similar trends to waveforms reported previously for cats walking overground and on a treadmill [25, 29, 41]. Additionally, hip abduction-adduction shown in Fig 5 in the present study similarly changed during swing (abduction increase followed by abduction decrease) as reported using a composite sagittal plane and frontal plane kinematic model [13]. This trend also was described in cats walking with either narrow or wide bases of support [13, 14].

The cat motor repertoire is extensive and includes sophisticated multidirectional movements beyond basic gait activities such as jumping, leaping, rolling, scratching and climbing. Capturing these movements and corresponding joint rotations will improve our understanding of motor control, and the multiplane model is well suited for evaluation of these movements in 3D. Furthermore, the multiplane model could be used to evaluate movements previously investigated using a sagittal plane approach. Additional richness of data (e.g. internal-external rotation) could be achieved for scenarios involving object negotiation [25], slope walking [29], or therapeutic intervention [26]. Multiplane models have been developed for humans [15, 18, 19, 36], and use of a multiplane model in cats will bolster applicability of the cat as a translational model to humans.

In the current study, hind limb kinematics trends were similar across healthy cats. It is important to note that the magnitudes of the kinematic outcomes, especially internal-external rotation and abduction-adduction, are dependent on the relative orientation of the defined anatomical coordinate systems based on segment motion capture markers applied to each cat. Therefore, subtle differences in defining the mediolateral and/or craniocaudal axes in each segment defining a joint may lead to kinematic outcomes that appear skewed. For instance limb internal-external rotation of the hip varied from 4.8° to 17.7° during gait for the cat reported in Fig 5. These data indicate that the hip is externally rotated throughout the step cycle. However, small differences between the kinematic model rotational axes and the cat hind limb physiological axes may lead to a small offset in magnitude and the definition of the neutral limb orientation (i.e., 0° orientation). Additionally, stifle flexion-extension at the start of stance ranged from 112° to 140° across the cats in this study, but stifle range of motion across cats was similar (40°). Therefore, it is important to use consistent marker placement across cats and evaluation time points to produce reliable data for assessment of multiplane kinematics trends. These best practice techniques are fundamentally necessary for a kinematic model and will improve likelihood of detecting differences across scenarios evaluated.

### Stifle prediction

Optical motion capture systems allow flexibility to assess gait across a range of tasks. However, it is known that skin motion around the stifle joint in cats and rats prevents reliable stifle tracking using skin-mounted markers and may lead to positional errors which could significantly affect kinematic calculations and lead to overestimation or underestimation of joint angles [31, 33]. Furthermore, the stifle instantaneous center may change as a function of flexion-extension angle as the tibia rolls around the femoral condyles [42]. Despite these concerns, optical motion capture systems that track skin-mounted markers are less invasive than surgically implanted tracking devices and less hazardous than x-ray cinematography. To minimize the influence of skin motion on stifle tracking, sagittal plane kinematic models in cats and rats have used mathematical projection techniques to estimate the stifle center based on distal tibial markers, the greater trochanter marker, and fibular and femoral lengths [25, 26, 30, 32] To our knowledge, direct comparison between x-ray cinematography, skin-mounted marker, and projection techniques have not been evaluated in the cat stifle. However, stifle joint projection techniques in rats showed better agreement with x-ray cinematography compared to skin mounted markers [31]. Peak differences in average craniocaudal and ventrodorsal stifle position during gait in rats using a skin-mounted stifle marker compared to a projected stifle marker based on measured segment lengths ranged from 5.0 – 7.5 mm while peak sagittal plane hip and stifle joint angles using these two techniques differed by as much as ± 20° [32]. Another study measured a peak difference in stifle flexion-extension angle during gait of 39 ± 6° between x-ray and skin-mounted marker techniques and 17 ± 11° between x-ray and projection techniques with peak errors occurring near paw contact [31].

In the current study, two stifle projection techniques were implemented and compared. In both techniques, the tibial mediolateral direction was defined based on the medial and lateral malleoli markers. A femoral mediolateral direction based on medial and lateral femoral condyle markers as done in canines [17, 20] and humans [18] was inappropriate for use in this study due to stifle skin motion artifact that occurs in the cat. Therefore, the femoral coordinate system was developed using the tibial mediolateral axis which couples the femoral and tibial segment orientations. This coupling between segments may influence hip and stifle internal-external rotation and abduction-adduction determined using the multiplane kinematic model. For instance, in the extended limb, internal rotation of the tibia relative to the femur would alter the femoral mediolateral axis causing it to be more internally rotated relative to the pelvis. This rotation may lead to underestimation of stifle internal rotation and overestimation of hip internal rotation. Therefore, this limitation should be considered when evaluating stifle and hip 3D rotations using the multiplane model. Despite this coupling, the multiplane model provides an in depth means to quantify rotations at each joint so that limb kinematic characteristics may be described in greater detail. Additional kinematic measures using the multiplane model could be developed to assess limb segment orientations that do not share a joint (e.g. tibia segment relative to pelvic segment).

### Sagittal plane and multiplane kinematics comparison

The sagittal plane and multiplane kinematic models predicted similar gait patterns within a single plane (Fig 6) which is similar to findings from a canine study evaluating 2D and 3D measurement systems [38]. However, in the present study, joint angle magnitudes differed by as much as 20° between the kinematic models, and each cat evaluated demonstrated differences across the joints analyzed. Subject variability is expected, and mean joint angles may differ normally by >10° across quadrupedal subjects [38]. Differences in hip joint flexion-extension occurred in the first cat, differences in tarsal joint flexion-extension occurred in the second cat, and differences in flexion-extension in the hip and tarsal joints occurred in the third cat. Differences between the sagittal plane and multiplane flexion-extension angles can be explained by the separation of internal-external rotation and abduction/adduction from flexion-extension in the multiplane model as these rotations are not isolated using the sagittal plane model and contribute to a composite flexion-extension. Additionally, joint angles were based on segment definitions from marker locations, and segment definitions may have differed between the sagittal plane and multiplane models.

Hip joint flexion-extension was consistently larger using the sagittal plane kinematic model compared to the multiplane kinematic model. This difference was not consistently present in a study evaluating dogs [38] and could be due to the use of separate 2D and 3D optical systems [38]. Furthermore, in the present study the ischial tuberosity and iliac crest were used to define the pelvis segment in the multiplane kinematic model while the iliac crest and greater trochanter were used in the sagittal plane model. This difference led to a larger flexion-extension angle in the sagittal plane model when the greater trochanter marker was more distal compared to the line connecting the iliac crest and ischial tuberosity markers used in the multiplane model (e.g. Cat 3 of Fig 6). In contrast, in Cat 2 of Fig 6, where the sagittal plane and multiplane kinematic models predicted similar sagittal plane hip joint angles, the greater trochanter marker was nearly in line with the iliac crest and ischial tuberosity markers.

Stifle flexion-extension was similar for the sagittal plane kinematic model and the multiplane kinematic model using both stifle projection techniques. These findings are consistent with findings reported for dogs [38]. This strong agreement also is due to the use of similar stifle projection techniques in both kinematic models. Both models predict a stifle marker based on the lateral malleolus marker, vector marker, greater trochanter marker, and fibula length. Additionally, the multiplane model stifle prediction techniques include femur length. As shown in Fig 4, each multiplane model stifle prediction technique led to similar stifle kinematics. In another study, triangulation of a knee marker in a rat kinematic model led to hip and knee flexion-extension angles with good agreement compared to bone-derived joint angles indicating knee marker prediction techniques are an effective approach when skin motion is significant [31].

Tarsal joint flexion-extension was consistently larger using the sagittal plane kinematic model compared to the multiplane kinematic model. This difference was not consistently present in a study evaluating dogs [38] and could be due to the use of separate 2D and 3D optical systems [38]. Furthermore, in the present study the lateral malleolus was used to define the tarsal segment in the sagittal plane kinematic model while the calcaneus marker was used in the multiplane model. This difference led to a smaller flexion-extension angle due to its more caudal position relative to the lateral malleolus. In Cat 1 of Fig 6, where the sagittal plane and multiplane kinematic models predicted similar sagittal plane tarsal joint angles, the calcaneus marker was nearly in line with the 5^th^ metatarsal and lateral malleolus markers. If the tarsal joint behaved as a hinge joint with the axis of rotation acting through the lateral malleolus, the lateral malleolus and calcaneus markers would both define the tarsus segment. However, the multiplane kinematic model predicted tarsal joint internal-external rotation and abduction-adduction indicating that the tarsal joint has more degrees of freedom than a hinge joint, which is consistent with tarsal joint passive range of motion measured in cats [43]. Therefore, the lateral malleolus and calcaneus markers may move independently of each other during ambulatory tasks which would influence measured tarsal kinematics.

Comparative data are reported for healthy cats in this study, but differences may be more prominent when comparing sagittal plane and multiplane kinematics in animals with abnormal gait patterns due to some sort of injury or developmental abnormality. Internal-external limb rotation may increase in pathologic gait [2, 39], and changes in internal-external rotation may modify the sagittal plane angle determined between three markers that define the joint. As the distal segment externally rotates relative to the proximal segment, the distal marker of the distal segment will rotate about the proximodistal axis in 3D space thereby changing the plane containing the three markers, the position of the distal marker relative to the proximal markers, and the angle between the markers. In this scenario no change in flexion-extension has occurred, but the angle between the markers has changed. The 3D kinematic model calculates angles between planes associated with three rotations [15, 17, 20] whereas the traditional sagittal plane model is constrained to calculating angles between intersecting lines within the same plane [20]. Therefore, changes in internal-external rotation while holding flexion-extension constant cannot be quantified except as a change in flexion-extension when using a sagittal plane kinematic model [20].

### 3D kinematics model validation

Joint angles determined using the 3D kinematic model based on markers placed on skin overlying bony landmarks were compared to joint angles determined using bone-oriented coordinate systems *ex vivo*. Joint angle patterns were similar across techniques but differed in magnitude due to slight offsets in orientation of the segment coordinate systems. After adjusting for the difference in orientation for each segment with the limb in a neutral position, measured joint angles differed between the two techniques by < 5° for all joints. These findings further support the use of stifle marker prediction techniques to define stifle kinematics in cats because differences between the bony landmark and marker triad coordinate systems were primarily due to differences in coordinate system setup alignment and not the use of a stifle marker prediction technique. External-internal rotation and abduction-adduction stifle and tarsal joint angles in the cadaver specimen were generally less in magnitude compared to *in vivo* joint angles. This difference may be due to motion of the hand not replicating actual gait that may have limited movements not contained within the sagittal plane. However, the movements applied to the cadaveric specimen were sufficient for testing and to show applicability of the multiplane model to determine flexion-extension, external-internal rotation, and abduction-adduction.

### Limitations

Overground walking is a daily activity in animals, and is therefore an important movement to evaluate in order to understand neurologic and biomechanical functions. Thus, these first assessments using a multiplane model were conducted on overground walking in healthy cats. Hind limb range of motion for other tasks or in injured cats may exceed the range of motion evaluated in this study. For instance, cats with spinal cord injury traversing pegs with predetermined spacing and contact area demonstrated greater hip flexion and knee extension when adopting a novel placing strategy in which a hind limb crosses the midline to place onto a contralaterally positioned peg [26]. Furthermore, for steps in which the ipsilateral limb misses a peg, the ipsilateral limb provides no body weight support, and the contralateral limb flexes to greater extent than while walking on a flat surface because the contralateral limb must support all pelvic region body weight throughout the majority of the step cycle [26]. Finally, negotiating an obstacle while walking necessitates increased vertical limb displacement for successful clearance [25]. Evaluation of other tasks requiring greater range of motion such as peg walking and obstacle negotiation using this 3D kinematic model is needed.

### Conclusions

The cat multiplane model developed in this study was used to determine flexion-extension, internal-external rotation, and abduction-adduction of the hip, stifle, and tarsal joints. The multiplane model brings greater interpretive power to gait assessment than sagittal plane assessment as it characterizes joint internal-external rotation and abduction-adduction in addition to flexion-extension. Three dimensional stifle kinematics were similar regardless of projection technique used, and although waveforms were similar, the absolute flexion-extension angles may differ when comparing the multiplane model to the sagittal plane model.

## Acknowledgements

The authors would like to thank Wilbur O’Steen for technical assistance.

